# Simple, rapid, and sensitive assay for the quantification of total polysaccharides to estimate extracellular polymeric substances (EPS) in soil

**DOI:** 10.1101/2025.05.22.654594

**Authors:** G. Bogar, J.T. Lennon, H. Vander Stel, S. Evans

## Abstract

Current methods for quantifying total polysaccharides in soil are expensive or require prolonged incubation of hazardous reagents at 100°C in glass vessels. We present a fast, sensitive, and inexpensive spectrophotometric assay quantifying polysaccharides in aqueous solution, featuring an instant reaction of room-temperature reagents using disposable tubes and cuvettes.

## Introduction

Soil microbes and plant roots influence soil physical (Benard et al., 2019; Blankinship et al., 2016; Bucka et al., 2021; Zheng et al., 2018) and chemical (Cotrufo et al., 2019) properties through the synthesis of mucilage and biofilms. These compounds, described as extracellular polymeric substances (EPS), bind strongly to clay minerals (Lin et al., 2016), and the rate of their deposition or loss may be important for long-term carbon sequestration in soil (Cotrufo et al., 2019; Schimel and Schaeffer, 2012; Sher et al., 2020). Soil EPS can be estimated by quantifying total polysaccharides (Lin et al., 2016) extracted with a gentle method that preserves and excludes intact cells (Redmile-Gordon et al., 2014). However, studies quantifying polysaccharides in soil often rely on a spectrophotometric assay developed nearly 70 years ago (DuBois et al., 1956) that is logistically challenging and not well-suited for modern microbial ecology laboratories.

## Methods, Results, and Discussion

We present a fast, sensitive, and inexpensive method to quantify total polysaccharides in aqueous solutions (Supplementary Material 1 – Verbose Protocol). Our method is based on previous work (Rasouli et al., 2014) that performed reactions in glass tubes, using 6.5% phenol and glucose/water solutions, incubated for 30 minutes. In our method, reactions are instead performed in 2.0 mL polypropylene swing-top tubes, using 5% phenol and glucose/1x PBS solutions, with an immediate reaction. We include additional steps and benchmarks to improve accuracy and ease of use for soil extracts. Data processing/analysis was conducted using R (R Core Team, 2024) and the ‘tidyverse’ package (Wickham et al., 2019).

Our method is more sensitive than the previous assay by a factor of 3.4, as estimated by slopes of standard curves (Rasouli et al., 2014, Figure 4a). However, as with the previous method, our results suggest that reaction-to-reaction variability can be high, and we recommend extremely clean materials and analytical duplicates or triplicates (Fig. 1). We also recommend that a new standard curve be calculated for each lot of reaction tubes, as we found variable baseline µg-scale reactable material between bags from suppliers, even with PCR-grade consumables. Furthermore, the order and rate of reagent combinations affects assay outcomes, and extreme caution must be taken due to the exothermic reaction of concentrated sulfuric acid and dilute phenol-sample solution (see Supplementary Material 1). We likewise advise extreme caution when attempting this protocol with materials and volumes other than those specified; it is possible that only the specific combination of a small reaction volume and thick polypropylene reaction tube prevents the tube from melting or cap from failing during the reaction.

**Fig. 1.**
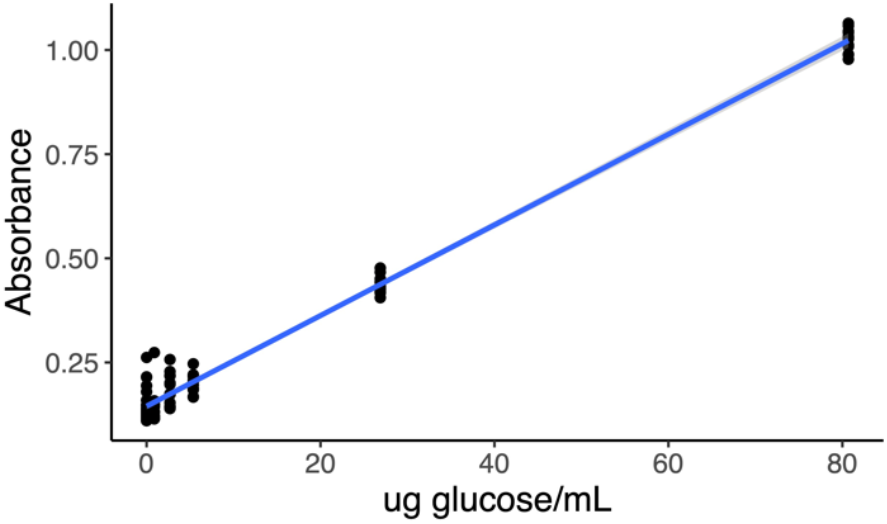
Total polysaccharide standard curve with 95% confidence interval, using the averages of analytical duplicate pairs: 27 blanks and 13 each of five glucose standards. ABS = 0.145 + 0.0109(µg glucose/mL). r^2^ = 0.99, Average standard deviation across curve (lower detection limit) = 2.8 µg glucose/mL. A new standard curve must be calculated for every new bag of reaction tubes.

To benchmark the effectiveness of our assay in soils, we extracted extracellular polysaccharides using the methods of Redmile-Gordon et al. (2014) from 68 field-moist agricultural soil samples (Jen Lau, *pers. comm.*), quantified glucose-equivalent polysaccharide per oven-dry mass of soil, and compared these with total soil carbon (Fig. 2). Soil was sampled from the top 20 cm of fields then growing corn (*Z. mays*), across three states of the U.S. Midwest from September – October 2021. Soils were transported and stored at 4°C until sieving, extraction, and quantification using this assay. We found that total extracellular polysaccharide concentrations ranged from 52 – 237 µg glucose-equivalent per gram oven-dry soil, with an average of 113 µg glucose-equivalent per g soil. This is approximately 1/3 the range and mean of water-extractable organic carbon (WEOC) from comparable soil samples (Boyer and Groffman, 1996).

**Fig. 2.**
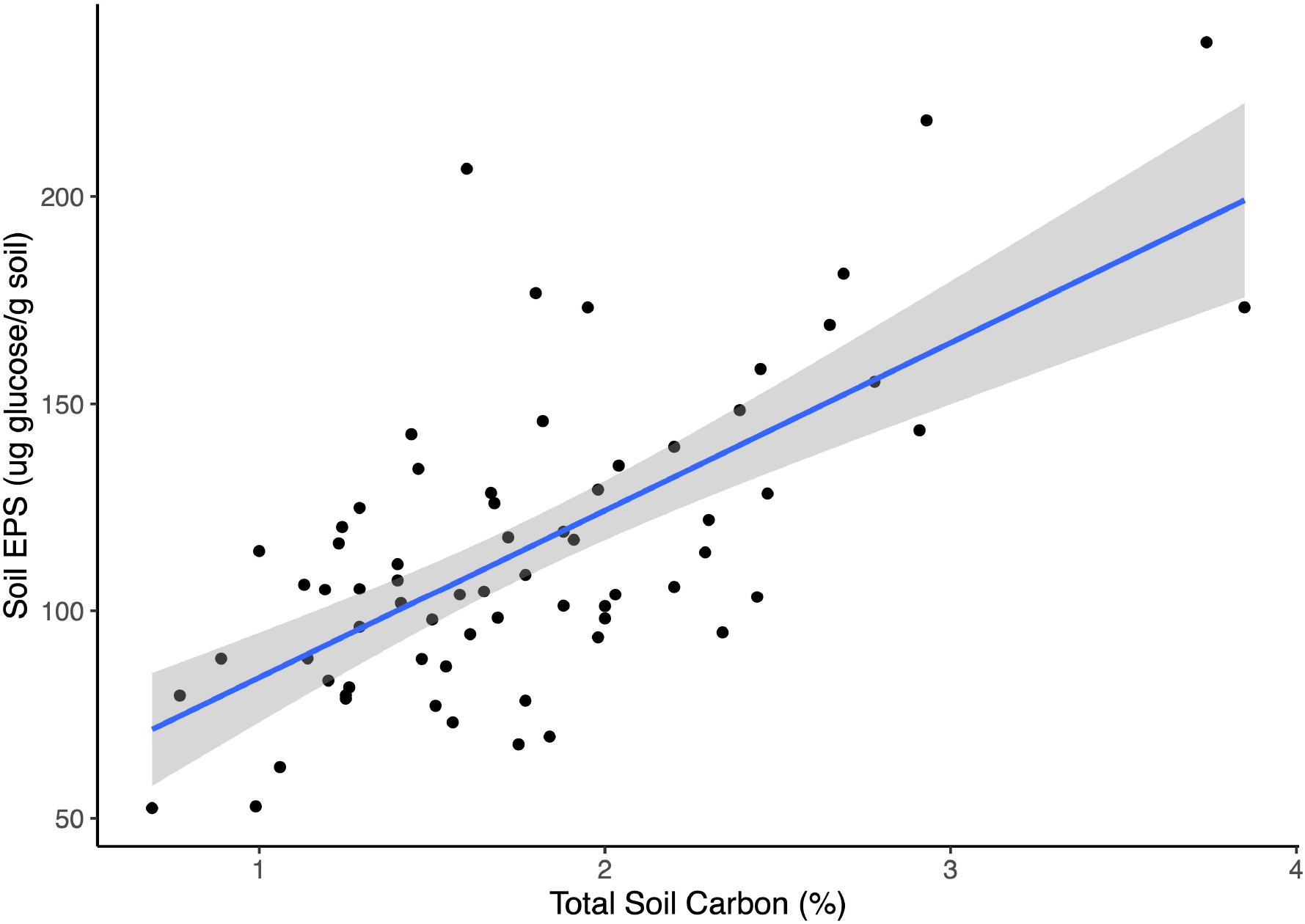
Linear regression of percent total soil carbon vs. total extracellular polysaccharides, with 95% confidence interval. F_66_ = 55.5, *P* = 2.5 * 10^−10^, r^2^ = 0.45.

There was a significant correlation (*r* = 0.67, *p* = 2.5 e-10) between %C and total extractable extracellular polysaccharides (Fig. 2), comparable to the strength of the relationship between %C and other commonly measured water-extractable C fractions, such as potassium permanganate oxidizable carbon (Culman et al., 2012).

We include several steps to improve accuracy and usability when measuring polysaccharides in soil extracts. First, we corrected for soluble soil pigments, which may not degrade during the dehydration reaction. Second, we tested and quantified the correlation of total extractable extracellular polysaccharides with ethanol-precipitated polysaccharides from which soluble sugars had been removed. When measuring polysaccharides as a proxy for large polymers (*e.g.,* microbial biofilm, root mucilage), extractions are generally precipitated to exclude small sugars. We extracted and quantified total and precipitated polysaccharides both in a mineral medium (81% sand, 19% clay) spiked with xanthan, and from soil in which corn (*Z. mays*) had grown, at the conclusion of a greenhouse drought stress experiment (Jen Lau, *pers. comm.*), to test the efficacy and requirement of polysaccharide precipitation. Solutions of total extractable polysaccharides were precipitated by adding ethanol to extractions to a final concentration of 80% at room temperature, followed by immediate centrifugation and resuspension in extraction solution (Alves et al., 2007; Brunchi et al., 2021).

Our precipitation recovered almost all large polysaccharides from extracts, but precipitation may not be necessary to estimate EPS in soil, given the strength of correlation we found between precipitated and total polysaccharides. We measured an average precipitation recovery of 92.5% from xanthan concentrations above 10 µg glucose equivalent/g soil (Supplementary Table S1), and a strong linear correlation between extracted and precipitated polysaccharides in soils (r = 0.98, Fig. 3). Further research is needed to determine the environmental contexts in which precipitation is required; it is likely still needed in circumstances where concentrations of soluble sugars are expected to vary between samples.

**Fig. 3.**
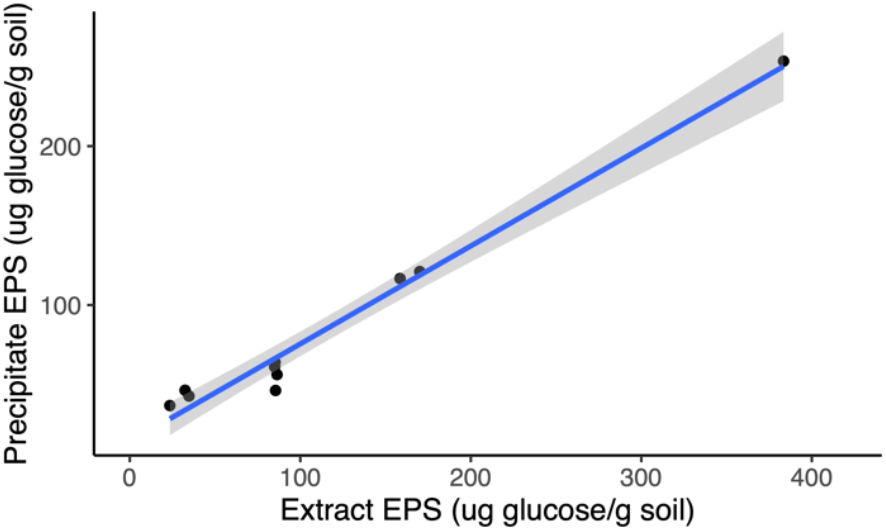
Linear regression of total extracted extracellular polysaccharide vs. precipitated extracellular polysaccharide with 95% confidence interval, extracted from soil in pots of corn (*Z. mays*) at the conclusion of a greenhouse drought stress experiment. F_8_ = 349, *P* = 7.0 * 10^−8^, r^2^ = 0.97. Precipitate EPS = 14.0 + 0.616(Extract EPS).

Our optimized and validated total polysaccharide assay should allow any lab with access to a flow hood and cuvette spectrophotometer to more easily study microbial biofilms and plant exudates (EPS) in soils and sediments. It may also be useful to researchers measuring non-structural carbohydrates (NSC) in plant tissues, or µg/mL-scale polysaccharides in any other cells or tissues.

## Supporting information

Supplementary Material 1 - Verbose Protocol

## Acknowledgements

This work was funded by NSF Award number 2009125 and 8685 “CNH2-L: Resilience to drought or a drought of resilience? The potential for interactions and feedbacks between human adaptation and ecological adaptation”. Thanks to Dr. Laura Bogar and the Dept. of Plant Biology at University of California, Davis, for laboratory equipment, resources, and helpful comments. Thanks to Dr. Jennifer Jones for preliminary assay validation.

## Authorship contribution statement

**Bogar, G.** Investigation, Methodology, Analysis, Original Draft, Review & Editing; **Lennon, J.T.** and **Vander Stel, H.** Review & Editing; **Evans, S.** Supervision, Methodology, Review & Editing

## Notes

### Competing Interest Statement

The authors have declared no competing interest.

