## Supplementary Material 1 - Verbose Protocol for "Simple, rapid, and sensitive assay for the quantification of total polysaccharides to estimate extracellular polymeric substances (EPS) in soil"

**EPS Quantification Protocol (modified from Rasouli et al., 2014)**

**Glade Bogar**

**EQUIPMENT**

-Personal protective equipment (PPE):

Lab coat

Resistant shoes

Long Pants

Splash-proof eye goggles

Disposable nitrile gloves

Mid-forearm-length butyl gloves

-Fume Hood

-Spectrophotometer (small enough to fit in fume hood, 490 nm read)

-1000 μL pipettor

-100-200 μL pipettor

-1000 mL glass beaker

-Vortexer

-PP (polypropylene) 15-mL tube racks (24-tube capacity)

-PP secondary containment for 15-mL tube racks

-Cuvette rack

-PP secondary containment w/lid, fitting cuvette rack

**REAGENTS**

-1000 μL concentrated sulfuric acid/sample (95-98% A300-212 Fisher Chemical)

-200 μL 5% phenol/sample (Phenol, 5% (w/v) Solution, 4.9-5.1%, Spectrum™)

-Glucose

0.18 g

-1X PBS supplies (2000 μL/sample):

8.00 g NaCl

0.20 g KCl

1.44 g Na_2_HPO_4_

0.24 g KH_2_PO_4_

OR

1X PBS pre-formulated mix

-Baking soda (sodium bicarbonate) for acid neutralization

**DISPOSABLE SUPPLIES**

-15.0-mL PP screw-top tubes (falcon) x24 – reusable during assay

-2.0-mL tubes (Eppendorf) – x2/sample, blank, standard

-Disposable ‘semi-micro’ (1.5 mL) PS (polystyrene) cuvettes – x2/blank, standard; x3 per sample

Note: these are commonly listed as “1.5-3.0 mL” cuvettes. In a spectrophotometer with an 8.5 mm read height, the actual minimum fill volume to accurately measure these cuvettes is ~500 μL regardless of their labeling. A volume of 0.7 mL fills these cuvettes to just below the top of the transparent read window, but the signal is read approximately halfway down this window.

-Disposable cuvette lids – x24 (re-usable if wanted, see hazardous waste protocol below)

-1000 μL pipette tips (up to 6/sample)

-100-200 μL pipette tips (1/sample)

-plastic transfer pipettes (2/sample)

-pH test strips

-quart Ziploc freezer bags for hazardous waste disposal (1 per 10 samples)

-PP containers with lids for dilute acid-contaminated tips

-screw-top glass FLPE (fluorinated high density polyethylene) bottles, at least 250 mL, for liquid waste

-HDPE (high density polyethylene) or PP buckets w/ tight-fitting lid for secondary storage of all ziploc bags containing hazardous waste.

**POTENTIAL HAZARDOUS WASTES PRODUCED**

**5% phenol**

NOTE: varying advice about correct storage conditions for aqueous phenol. Phenol concentration and acceptable damage to storage material (discoloration, swelling, weakening, surface dissolution) may account for variability. It seems likely that for 5% phenol, at room temperature, for a relatively short time, PP is sufficient. However, for high concentrations and long-term storage and disposal, fluorinated polyethylene (FLPE) or glass is recommended.

IF NEEDED: Storage: 50 mL falcon tube (sealed) within a PP container (w/lid, taped shut). Waste tag on PP container. Waste should not leave fume hood until collected.

**5% phenol-contaminated tips**

Storage: Ziploc freezer bag within HDPE or PP secondary containment bucket w/lid. Waste should not leave fume hood until collected.

**Concentrated sulfuric acid (98%)**

NOTE: If diluted to at least 50% (preferably to 10%, ~2M) long-term storage in PP is acceptable.

NOTE: If diluted with water, concentrated sulfuric acid should be diluted to a maximum final concentration of 10% to avoid excessive heating. Sulfuric acid will become dangerously hot if diluted to more than 30%, and may boil if diluted to more than 40% final concentration.

Mitigation: Make a ~1M solution of sulfuric acid by adding 20x the volume of water (compared to acid) to a glass container with a screw-top lid, then adding the acid to this container, and dispose as hazardous waste.

**Concentrated sulfuric-acid-contaminated tips**

NOTE: concentrated sulfuric acid will degrade PP, acid must be diluted before storage and disposal as hazardous waste.

Mitigation: Fill 1000-mL glass beaker to ~500 mL with water. Before ejecting pipette tips that have been used to transfer concentrated sulfuric acid, rinse by pipetting twice up-and-down in water. Eject rinsed pipette tips into storage container.

After quantification, place beaker with dilute acid solution in lab sink. Add 10g sodium bicarbonate to dilute acid solution. Stir to react. Test with pH strip. If pH < 6.0, repeat. When pH > 6.0, pour remaining sodium sulfate solution down lab sink.

Storage: PP container w/lid, taped shut.

**70% sulfuric acid/1% phenol-contaminated transfer pipettes**

Place in a Ziploc quart freezer bag. Store sealed freezer bags in PP or HDPE bucket w/tight-fitting lid. Waste should not leave fume hood until collected.

**Phenol- and acid-contaminated 2.0-mL tubes, sealed**

Place in a Ziploc quart freezer bag. Store sealed freezer bags in PP or HDPE bucket w/tight-fitting lid. Waste should not leave fume hood until collected.

**Phenol- and acid-contaminated cuvettes + 70% sulfuric acid, 1% phenol solution**

Carefully remove cuvette lid, and pour 70% sulfuric acid, 1% phenol solution into a screw-top FLPE or glass bottle. Place empty cuvettes in a Ziploc quart freezer bag. Store sealed freezer bags in PP or HDPE bucket w/tight-fitting lid. Waste should not leave fume hood until collected.

Alternatively, place sealed cuvette, with solution inside, into a large screw-top glass or PP container for greater security. Note that this will take up a lot of space, about 0.5 L per set of 12 samples.

**PREPARATION**

**Make 1L 1X PBS solution:**

Rinse two 1000 mL autoclavable bottles three times with nanopure water.

Add to one 1000-mL autoclavable bottle:

8.00 g NaCl

0.20 g KCl

1.44 g Na_2_HPO_4_

0.24 g KH_2_PO_4_

990 mL nanopure water

Add a magnetic stir bar, and stir until completely dissolved.

When dissolved, (~10 minutes) pour ½ of 1X PBS soln into second 1000-mL autoclavable bottle.

Autoclave on 0.5 L, liquid setting (40 minutes 121 degrees C, slow depressurize)

Allow to cool.

**Make glucose standard solutions:**

Label six 15-mL falcon tubes:

EPS BLANK

EPS STD #1

EPS STD #2

EPS STD #3

EPS STD #4

EPS STD #5

Label three 2.0 mL Eppendorf tubes:

EPS STD 1000x STOCK

EPS STD 100x STOCK

EPS STD 10x STOCK

Make a 1000x std stock soln.:

Weigh 0.18g glucose directly into “EPS STD 1000x STOCK” tube.

Add 1000 μL 1x PBS.

Vortex thoroughly to dissolve.

Concentration: ~160,000 ug/mL glucose

Note the effect of adding 0.18g * (1mL/1.56g) = 0.1154 mL glucose; total final volume = 1115.4 μL

Make a 100x std stock soln:

Add 900 μL 1X PBS to “EPS STD 100x STOCK” tube.

Add 100 μL 1000x std stock soln.

Vortex to mix.

Make a 10x std stock soln:

Add 900 μL 1X PBS to “EPS STD 10x STOCK” tube.

Add 100 μL 100x std stock soln.

Vortex to mix.

Make EPS Std #5:

Add 4750 μL 1x PBS to “EPS STD #5” tube.

Add 250 μL 10x std stock soln.

Vortex to mix.

Final concentration: 80.69 ug/mL glucose.

Make serial dilutions of Std #5 to make Stds 4-1:

Add 3335 μL 1X PBS to each of tubes labeled Std #4.

Add 4800 μL 1X PBS to each of tubes labeled Std #3.

Add 2500 μL 1X PBS to each of tubes labeled Std #2.

Add 3335 μL 1X PBS to each of tubes labeled Std #1.

Add 1665 μL Std #5 to Std #4, vortex (26.87 ug/mL glucose).

Add 1200 μL Std #4 to Std #3, vortex (5.37 ug/mL glucose).

Add 2500 μL Std #3 to Std #2, vortex (2.69 ug/mL glucose).

Add 1665 μL Std #2 to Std #1, vortex (0.89 ug/mL glucose).

Add 10 mL 1x PBS to BLANK tube.

**PROTOCOL**

THIS PROTOCOL INVOLVES CHEMICAL AND PHYSICAL HAZARDS INCLUDING INSTANT HEATING OF A SOLUTION OF PHENOL AND CONCENTRATED SULFURIC ACID FROM ~20 °C TO ~100+ °C.

A TECHNICIAN WITHOUT CORRECT PPE WILL BE INJURED WHEN FOLLOWING THIS PROTOCOL.

READ CONCENTRATED SULFURIC ACID AND PHENOL MSDS.

ALWAYS WORK WITH PHENOL AND CONCENTRATED ACID IN A FUME HOOD.

ALWAYS WEAR RESISTANT SHOES, LONG PANTS, LAB COAT, SPLASH GOGGLES, APPROPRIATE GLOVES WHEN HANDLING CONCENTRATED SULFURIC ACID AND PHENOL.

IF DILUTING ACID: ALWAYS ADD ACID TO WATER, NEVER WATER TO ACID – 10% MAXIMUM FINAL CONCENTRATION.

**Notes**

This method has been developed to measure total sugars and polysaccharides in a soil extraction (As described in Redmile-Gordon et al., 2014, ‘Measuring the soil-microbial interface: Extraction of extracellular polymeric substances (EPS) from soil biofilms’) using a cuvette spectrophotometer. When concentrated sulfuric acid is added to a mixture of dilute polysaccharides and phenol, a violent exothermic reaction dehydrates and condenses polysaccharide into furfural, which absorbs light strongly at 490 nm. This method can detect total polysaccharide concentrations between ~2.5 – 180 ug/mL glucose equivalent. Using the reaction volumes described, with the specified materials and methods, the assay solution heats rapidly from ~20 °C to ~100 °C but does not boil, does not destroy the tubes, and does not evolve enough heated gas to generate pressure within sealed tubes. Technicians should use EXTREME CAUTION when modifying or adapting this protocol. Changes could make this protocol more dangerous.

This method has been optimized for sensitivity by changing solution proportions (Rasouli et al., 2014, ‘Characterization and improvement of phenol-sulfuric acid microassay for glucose-based glycogen’). Sensitivity could be further increased by changing solution temperatures and incubation times. Taylor, 1995 (‘A modification of the phenol/sulfuric acid assay for total carbohydrates giving more comparable absorbance’) showed that assay sensitivity is increased by heating the phenol/sugar solution to 100 °C prior to reaction with sulfuric acid. However, when phenol solution begins significantly above ambient temperature, the additional exothermic heat instantly boils the solution.

An additional novel control needed for soil extracts is the 490nm absorbance of organic acids and pigments extracted along with extracellular polysaccharides. This is controlled by measuring a dilution of each soil extract without reaction first, and subtracting this absorbance from the final reacted absorbance.

Analytical replicate variability can be high. For this reason, all absorbance values should be calculated from averages of (at least) analytical duplicates. Ideally, triplicates or even quadruplicates are recommended.

Sets of 24 reactions maximum are recommended to allow adequate technician focus while pipetting and handling concentrated acid. So:

When running an initial standard curve:

Reaction blank (x4) + Standards #1-5, (x4)

= 24 reactions total

If running a full standard curve in every set:

Reaction blank (x2) + Standards #1-5, (x2) + Samples, x6 (x2)

= 24 reactions total

If running positive and negative controls only (blank and std #5 - recommended):

Reaction blank (x2) + Standard #5, (x2) + Samples, x10 (x2)

= 24 reactions total

**BEGIN**

**Preliminary Test: Standard Curve and Analytical Variability**

Label 2.0 mL PP tubes: x4 per blank and standard.

Label cuvettes (outside light path): x2 per blank and standard. Place in rack.

Place plastic transfer pipettes (x24) in an easy-to-reach configuration, such as a clean beaker, bulb-side up.

Place x24 15 mL tubes (without lids) into PP tube racks; place tube racks into PP secondary containment.

Explanation: 15 mL tubes hold 2.0 mL tubes into which acid is added to phenol/sample solution, for safety and heat dissipation. See Figure 1.

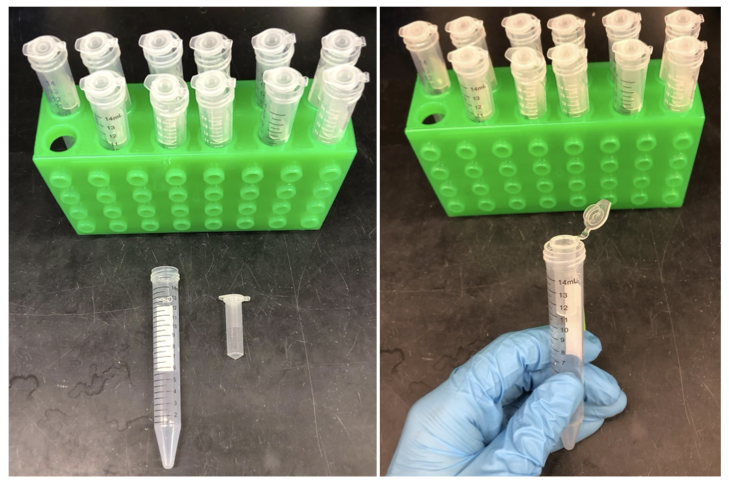

**Figure 1. Demonstration of tube arrangement for polysaccharide assay.**

Prepare hazardous waste receptacles:

·	Fill a 1000 mL beaker to ~500 mL with tapwater, label “dilute acid”, and move to fume hood.

·	Label a PP container “rinsed tips – non-hazardous”, move to fume hood.

·	Label 1 quart-size Ziploc freezer bag: “70% concentrated sulfuric acid/29% 1X PBS/1% phenol”, and move to fume hood.

·	Label 1 large bucket with a tight-fitting lid: “Secondary containment – Ziploc bags filled with plastic contaminated with 70% concentrated sulfuric acid/29% 1X PBS/1% phenol”, and move to fume hood. Attach an appropriate waste tag for pickup.

·	Label a 250+mL FLPE or glass bottle “70% concentrated sulfuric acid/29% 1X PBS/1% phenol”, and move to fume hood. Attach an appropriate waste tag for pickup.

Pipette 100 μL of each blank or standard into each of x4 labeled 2.0 mL tubes.

Wear all required PPE for phenol and concentrated sulfuric acid:

·	Forearm-length butyl or viton-butyl gloves, over standard nitrile gloves.

o	Viton-butyl gloves are ideal, but as the phenol concentration is low, thick butyl gloves are acceptable. 5% phenol solution will react with butyl.

·	Lab coat.

·	Splash goggles.

·	Face shield (if pouring concentrated sulfuric acid i.e., into a smaller container for pipetting).

·	Recommended: Face mask to protect nose and mouth.

Move to hood:

·	Vortexer

·	1000-μL pipettor and pipette tips

·	Hazardous waste receptacles

·	5% phenol solution

·	Concentrated sulfuric acid

·	15-mL tubes x24, in tube racks, in secondary PP container

·	Labeled cuvettes, x24, in rack

·	Cuvette lids

·	Labeled 2.0 mL tubes, with 100 μL blank/standard

Place each 2.0 mL tube into one of the 15 mL tubes. 2.0 mL tubes should rest within the open 15-mL tubes when open or closed, so that the 2.0 mL tube can be picked up, opened, and closed without touching its bottom or sides. 15-mL tubes also serve as secondary containment for each individual 2.0 mL tube (Figure 1).

Using a 1000-μL pipette, add 100 μL 5% phenol to each 2.0 mL tube and close 2.0 mL tube lid tightly. Dispose of contaminated tips in Ziploc freezer bag; seal bag.

Note: 1000 μL pipette tips were used as their longer size ensured that the pipettor would never contact the inside of the bottle containing 5% phenol solution.

Vortex all tubes to mix, replace in 15 mL tubes.

Mixing technique and reaction time makes a significant difference in the results for the next step; it is critical to mix the acid/phenol solution briskly. Try to repeat these steps exactly between reactions.

Starting with one set of blank/standards, as quickly as safely possible:

·	Pick up each 15 mL tube containing a 2.0 mL tube, itself containing the reaction solution (phenol/sample).

·	Open the 2.0 mL tube slowly and carefully, minimizing the chance of droplets on the lid getting on the gloves or flying out of the tube.

·	While holding the 15 mL tube, **rapidly** pipette 500 μL of concentrated sulfuric acid **directly** into the phenol/sample mixture at the bottom of the 2.0 mL tube. Do not pipette dropwise nor onto the inner sides of the 2.0 mL tube.

o	Depending on pipetting velocity and ambient temperature, solution may boil. **Use extreme caution.**

·	Quickly rinse the pipette tip by submerging in 500 mL tapwater and pipetting up and down twice.

·	Eject pipette tip into “rinsed tips – non-hazardous” container.

·	Seal the 2.0 mL tube containing acid/phenol solution tightly.

·	Pick up the **sealed** 2.0 mL tube by the lid hinge.

·	Gently shake 2.0 mL tube 10 times to mix, taking care to incorporate all droplets of phenol solution on sides or lid of tube. However you do it, try to shake all tubes in exactly the same way, for the same amount of time.

o	**At this step, solution heats to 100+** °**C.** Using Olympus Plastics brand 2.0 mL PP hinge-top tubes, heated gas pressure did not pop open the tube, nor did the tube become uncomfortable to touch through thick butyl gloves. However, other types of tube did pop open slightly on shaking, and tubes may be too hot to handle with thinner gloves.

·	Replace sealed 2.0 mL tube in 15 mL tube to cool down, and move to the next reaction.

When you have run through one set of blank and standards 1-5, move on to the next set, and the following two after that. Reaction tubes left overnight had much higher absorbance than those read immediately after the reaction, so the absorbance does change over time, but this was not apparent during the timescale of a single set of reactions (~20 minutes). To be safe, analytical duplicates are run in blocks.

By the time the final reaction is complete, the first tube should have reached ambient temperature. If so, carefully transfer the entire volume of solution (~700 μL) of each tube into the corresponding cuvette, using the transfer pipettes.

Dispose transfer pipettes in Ziploc bag, and seal.

Seal cuvettes with cuvette lids.

Repeat transfers in reaction order.

Place cuvette rack into PP box with a tight-fitting lid.

Remove Butyl/Viton-Butyl glove on your dominant hand.

Do not touch door handles/spectrophotometer with potentially contaminated gloves.

Move cuvette rack to fume hood containing spectrophotometer.

Read and record cuvette absorbances at 490 nm:

·	Set ‘Blank’ with no cuvette in the spectrophotometer.

·	Read each Blank, Standard sample twice

o	Recording each cuvette twice to check for spectrophotometer issues.

Replace cuvette rack in PP box and seal with lid.

Move back to first fume hood.

Replace butyl/viton-butyl glove on dominant hand.

Either:

One at a time, carefully remove cuvette lids and pour contents into 250+mL FLPE or glass bottle labeled “70% concentrated sulfuric acid/29% 1X PBS/1% phenol”.

Provided cuvette lids never touched cuvette contents, lids can be re-used; if so, store in a ziploc bag labeled “Cuvette lids – contaminated with 70% concentrated sulfuric acid/29% 1X PBS/1% phenol” in fume hood.

Place empty cuvettes into Ziploc freezer bag labeled “70% concentrated sulfuric acid/29% 1X PBS/1% phenol”. Seal Ziploc bag.

Or:

Do not unseal cuvettes, place sealed cuvettes directly into 250+mL FLPE or glass bottle labeled “70% concentrated sulfuric acid/29% 1X PBS/1% phenol”. This minimizes handling of 70% concentrated sulfuric acid/29% 1X PBS/1% phenol solution, but requires an equal number of cuvette lids to cuvettes, and requires a lot of space. This is recommended if you already have access to very large screw-top glass or PP containers for hazardous waste disposal.

Place sealed Ziploc freezer bag into large bucket with a tight-fitting lid, labeled “Secondary containment – Ziploc bags filled with plastic contaminated with 70% concentrated sulfuric acid/29% 1X PBS/1% phenol”. Seal bucket.

Neutralize dilute sulfuric acid solution created by rinsing pipette tips:

a. Remove “dilute acid” beaker from fume hood, place in lab sink.

b. Add 10g sodium bicarbonate to dilute acid beaker, stir to mix.

c. When CO2 creation stops, measure pH of resulting sodium sulfate solution with pH strip.

a. If pH > 6, rinse solution down drain.

b. If pH < 6, repeat steps (b) and (c) until pH > 6, then rinse down drain

c. Note that it takes 28 grams sodium bicarbonate to neutralize 1 mL concentrated sulfuric acid.

If hood will be used for other work, spray down hood interior with 1% baking soda solution and wipe down to neutralize trace acid.

Average multiple absorbances for each cuvette. Average analytical duplicates to produce two absorbance values for each blank and standard.

Evaluate standard absorbances:

·	Blank: Average 0.145, Range +/- 0.080; Std Dev = 0.038

·	Std 1: Average 0.149, Range +/- 0.080; Std Dev = 0.038

·	Std 2: Average 0.186, Range +/- 0.059; Std Dev = 0.037

·	Std 3: Average 0.200, Range +/- 0.040; Std Dev = 0.022

·	Std 4: Average 0.436, Range +/- 0.036; Std Dev = 0.020

·	Std 5: Average 1.023, Range +/- 0.043; Std Dev = 0.027

If the range for each standard is larger than these values, it is likely that there is a problem with mixing after acid addition. If the absorbance average for each standard is higher than these values, it is likely that there is some baseline contamination in the tubes.

**Sample Assay**

Protocol for samples is identical to above, with the added measurement of unreacted absorbance from soil extracts. Add the following steps into the protocol above.

After Line 308 - Label pigment cuvettes: x1 per sample.

After line 340:

Pipette 100 μL of each sample into each of labeled pigment cuvettes.

Add 600 μL 1X PBS to each of x1 labeled cuvettes in second cuvette rack, pipetting up and down to mix with 100 μL soil extract.

Move cuvette rack to fume hood containing spectrophotometer.

Read and record cuvette absorbances at 490 nm:

·	Set ‘Blank’ with no cuvette in the spectrophotometer.

·	Read each sample twice

o	Recording each cuvette twice to check for spectrophotometer issues.

Discard pigment cuvettes as non-hazardous waste.

These absorbance values allow for correction for soil pigments, many of which absorb at 490 nm.
